# DNA replication during acute MEK inhibition drives acquisition of resistance through amplification of the *BRAF* oncogene

**DOI:** 10.1101/2021.03.23.436572

**Authors:** Prasanna Channathodiyil, Anne Segonds-Pichon, Paul D. Smith, Simon J. Cook, Jonathan Houseley

## Abstract

Mutations and gene amplifications that confer drug resistance emerge frequently during chemotherapy, but their mechanism and timing is poorly understood. Here, we investigate *BRAF*^*V600E*^ amplification events that underlie resistance to the MEK inhibitor selumetinib (AZD6244/ARRY-142886) in COLO205 cells, a well-characterised model for reproducible emergence of drug resistance, and show that *de novo* amplification of *BRAF* is the primary path to resistance irrespective of pre-existing amplifications. Selumetinib causes long-term G1 arrest accompanied by reduced expression of DNA replication and repair genes, but cells stochastically re-enter the cell cycle during treatment despite continued repression of pERK1/2. Most DNA replication and repair genes are re-expressed as cells enter S and G2, however, mRNAs encoding a subset of factors important for error-free replication and chromosome segregation including TIPIN, PLK2 and PLK3 remain at low abundance. This suggests that DNA replication in drug is more error prone and provides an explanation for the DNA damage observed under long-term RAF-MEK-ERK1/2 pathway inhibition. To test the hypothesis that DNA replication in drug promotes *de novo BRAF* amplification, we exploited the combination of palbociclib and selumetinib. Combined treatment with selumetinib and a dose of palbociclib sufficient to reinforce G1 arrest in selumetinib-sensitive cells, but not to impair proliferation of resistant cells, delays the emergence of resistant colonies, meaning that escape from G1 arrest is critical in the formation of resistant clones. Our findings demonstrate that acquisition of MEK inhibitor resistance often occurs through *de novo* gene amplification and can be suppressed by impeding cell cycle entry in drug.

## Introduction

The development of targeted anti-cancer drugs has improved treatment efficacy and reduced side effects but drug resistance still limits long-term patient survival (1, 2). Mutations and gene amplifications affecting the drug target or proteins in downstream pathways allow re-emergence of tumours that are refractory to treatment with the original and related chemotherapeutics (3, 4).

Constitutive activation of the RAS-RAF-MEK-ERK1/2 pathway (hereafter, ERK1/2 pathway), resulting from mutational activation of BRAF or KRAS proteins, occurs in the majority of melanomas and colorectal cancers (5, 6). Consequently, the ERK1/2 pathway is a major target for drug development and inhibitors of RAF and MEK are approved for treatment of melanoma, whilst ERK1/2 inhibitors are undergoing clinical trials; however, patients often relapse with drug-resistant tumours (7, 8). For example, Selumetinib (AZD6244 / ARRY-142886) is a highly specific MEK inhibitor (MEKi) that suppresses constitutive activity of the ERK1/2 pathway, shows promise in pre-clinical studies (9, 10), but resistance to MEKi often arises through amplification of *BRAF* or *KRAS* (11-15).

Cancer cells are genetically heterogeneous and rare pre-existing mutations that confer drug resistance may be positively selected under drug treatment (16, 17). However, *de novo* mutations that occur during drug exposure can also cause resistance (18), in which case cells must survive initial drug application and then acquire mutations that restore proliferation. In culture, and recently *in vivo*, small numbers of drug tolerant persister (DTP) cells have been observed to survive extended treatment with targeted chemotherapeutics (19-22). DTPs exist in a non-proliferative or slow cycling state with gene expression patterns and metabolic states distinct from untreated and resistant populations (19-21, 23-26). However, proliferative colonies routinely emerge from DTPs in the presence of drug after long periods of apparent stasis, marking the DTP state as a precursor to resistance (18, 20-22). DTPs do not stem from a genetically defined subpopulation in the parental cell line and are not inherently drug resistant since removal from drug restores normal susceptibility (19, 21, 22), but colonies of resistant cells derived from DTPs carry drug resistance mutations of unknown provenance and emerge with kinetics consistent with *de novo* mutation (18, 27). Recently, inhibition of EGFR, which acts upstream of the ERK1/2 pathway, was shown to downregulate DNA replication and repair genes while inducing error prone DNA polymerase genes, which may indicate entry to a mutagenic state in response to drug exposure (28-30).

The cause of *de novo* mutation in DTPs is of great interest as mutagenic mechanisms that act during therapy could be inhibited to slow the acquisition of resistance. Here, we have made use of COLO205 cells treated with the MEKi selumetinib; this is a well-established and reproducible model of tolerance converting to resistance through gene amplification of the addicted oncogene, *BRAF*^*V600E*^ (12,13). We demonstrate that MEKi resistance arises predominantly through *de novo BRAF* amplifications in colorectal cancer cells. Although expression of DNA replication and repair genes is decreased during treatment as previously reported, we find that most are re-expressed as DTPs sporadically enter S phase. However, expression of a subset of genes important for error-free DNA replication remains low throughout the cell cycle in drug, and reducing the frequency of DNA replication events in drug delays the formation of selumetinib-resistant clones. Our results implicate DNA replication in drug as a major driver of *de novo* mutation leading to drug resistance in DTPs.

## Results

### Selumetinib resistance in COLO205 cells arises primarily through de novo BRAF^V600E^ amplification

Colorectal cancer cells carrying the *BRAF*^*V600E*^ mutation can overcome MEK inhibition by amplification of *BRAF*^*V600E*^, increasing levels of BRAF^V600E^ protein to activate more MEK and sustain ERK1/2 activity (11, 12). However, such *BRAF*^*V600E*^*-*amplified cells become addicted to MEKi; withdrawal of MEKi drives excessive MEK-ERK1/2 activity due to an over-abundance of BRAF^V600E^, resulting in cell cycle arrest, senescence and apoptosis (12, 31). Since a level of *BRAF*^*V600E*^ amplification that is sufficient for resistance should not be tolerated in a drug-naїve cell, *de novo BRAF* amplifications formed during treatment seem likely to underlie MEKi resistance. However, pre-existing *BRAF*-amplified cells (∼4%) have been reported in drug naïve colorectal cancer cells (11). These conflicting observations led us to investigate the contributions of pre-existing and *de novo BRAF*^*V600E*^ amplifications to the emergence of MEKi resistance in colorectal cancer cell lines.

Copy number profiling of 7 selumetinib-resistant COLO205 clones, derived from 7 independent drug treated cultures, revealed 3 (Resistant e, f & g) sharing identical *BRAF* amplifications that must have been present in the parental cell line prior to drug exposure (Fig. 1A, B). However, the other 4 resistant lines (Resistant a-d) carried unique amplicon structures that either formed *de novo* or emerged from a highly heterogeneous *BRAF*-amplified population in the parental line. To separate these possibilities, we erased existing population heterogeneity by deriving 10 clonal COLO205 cell lines, all of which were sensitive to selumetinib and generated resistant clones by prolonged drug exposure. Copy number profiles of 8 resistant cell lines derived independently from 3 of the clonal COLO205 cell lines revealed 7 unique *BRAF* amplicons representing different *de novo* amplification events, and one resistant line with no detectable amplification that must have gained resistance by a different mechanism (Fig. 1C). Importantly, the time taken for resistant colony formation in 9 of the clonal cell lines was identical to parental COLO205 cells, with the other line being only slightly delayed (Fig. 1D). This shows that *BRAF* amplification occurs frequently in COLO205 cells and that pre-existing *BRAF*-amplifications in the parental COLO205 cell line contribute little to the timing of selumetinib resistance.

**Figure 1:**
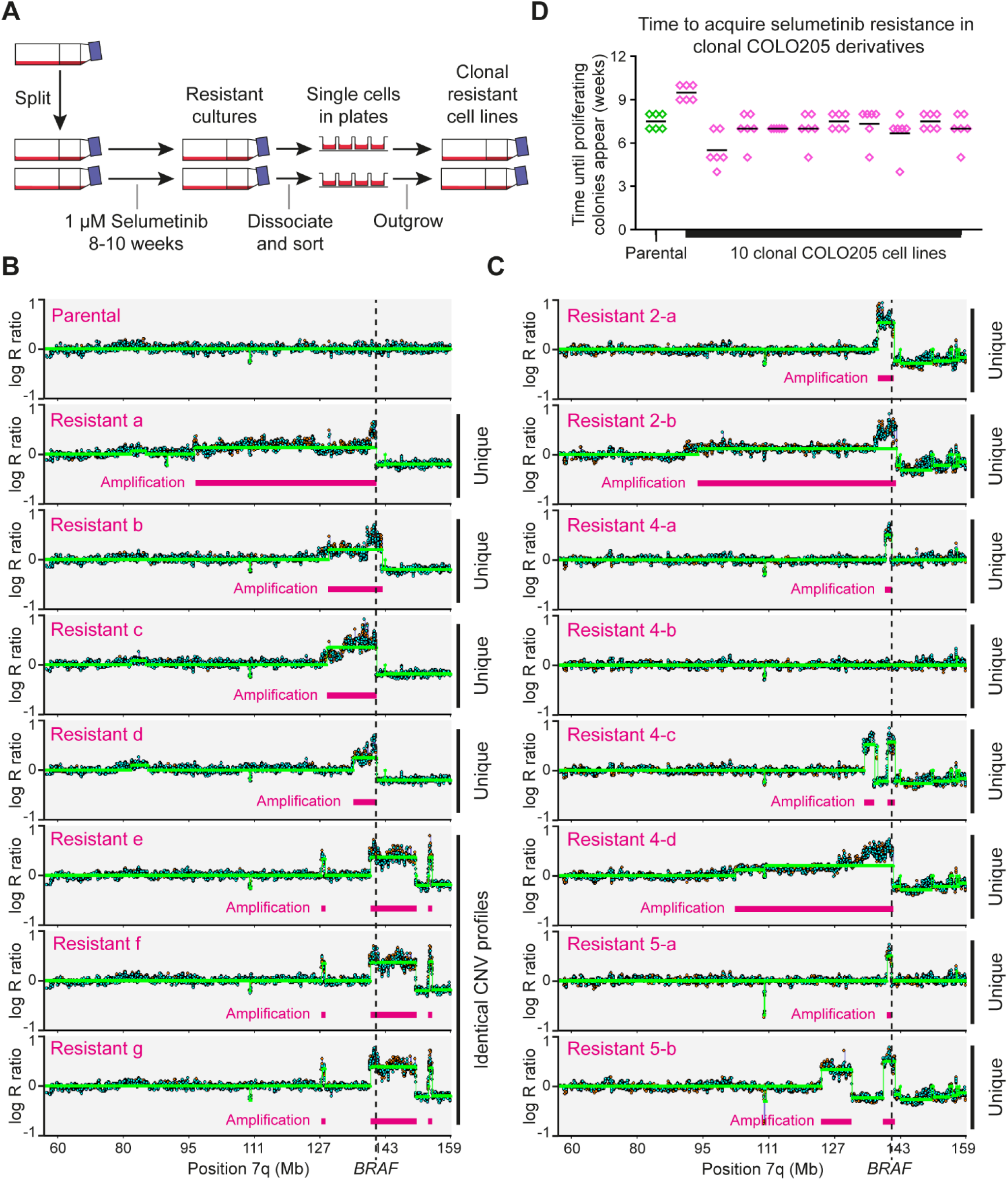
Contributions of pre-existing and *de novo* gene amplifications to the emergence of selumetinib resistance in COLO205 cells. **A**: Experimental design for analysing reproducibility of resistance. COLO205 cells were cultured in 24 individual 25 cm^2^ cell culture flasks in media containing 1 µM selumetinib. Media and drug were changed weekly until colonies of proliferating cells were observed, at which point single cells were isolated by flow cytometry and expanded into separate drug resistant cell lines in the presence of 1 µM selumetinib. **B**: Copy number profiles of the *BRAF* locus in parental COLO205 cells and 7 selumetinib resistant cell lines, determined using CytoSNP 850K BeadChip arrays (Illumina). Log_2_ ratio plots of copy number for the q arm of chromosome 7 are shown, including the location of *BRAF* (dotted line) and amplified regions (pink bars). Each resistant cell line derived from clonal amplification of individual selumetinib resistant cells from independent drug treatment flasks, but note that cell lines e, f and g show identical and highly characteristic copy number amplification profiles. **C**: CNV profiles of the *BRAF* locus in 8 selumetinib resistant lines obtained from clonal parental cell lines, each resistant cell line derived by clonal amplification from an independent drug treatment flask. Three clonal parental cell lines (clonal cell lines 2, 4 and 5) were used, with 4 resistant clones derived from cell line 4 and 2 each from cell lines 2 and 5. Note that all CNV profiles are different, and that cell line 4-b has become resistant without amplification of the *BRAF* locus or any region detectable by array-based CNV analysis. **D:** Time taken for proliferating selumetinib resistant clones to emerge from parental and 10 different single cell-derived COLO205 cell lines. For each cell line, cells were seeded in 6-well plates and treated individually with 1 µM selumetinib after 24 hours. Media and drug were changed weekly, and the time taken to colony formation was recorded (in weeks). The time to resistance was not significantly different between the parental line and any of the clonal lines (p>0.5) by a Kruskal-Wallis test.

### Individual cells enter the cell cycle even under acute MEK inhibition

Inactivation of the ERK1/2 pathway by MEK inhibition induces G1 cell cycle arrest in *BRAF*^*V600E*^ cell lines including COLO205 (32, 33). The treatment conditions used here result in growth arrest of COLO205 cells with no passaging required across six or more weeks in the presence of drug; instead, gradual cell death occurs over many weeks with remaining DTPs aggregating into large bodies from which resistant colonies often, but not always, emerge (Fig. 2A). Since *de novo* gene amplification normally occurs through errors in DNA replication or chromosome segregation (34, 35), we assessed whether selumetinib-treated cells escape G1 arrest using incorporation assays for the thymidine analogue ethynyl deoxyuridine (EdU). A 4-hour EdU pulse applied 24 hours after addition of selumetinib to COLO205 cells labelled 4.9 ± 0.3% of cells, compared to 43 ± 4% of control cells (Fig. 2B), confirming that a fraction of cells undergo DNA replication even in the presence of selumetinib. Equivalent results were obtained in clonal COLO205 cell lines (Fig. 2B), and EdU positive cells were detectable at all times analysed up to at least 7 days after selumetinib application (Fig. S1A). Therefore, COLO205 cells occasionally enter the cell cycle and initiate DNA replication during extended selumetinib treatment despite the seemingly robust G1 arrest. To ensure that escape from arrest is not unique to COLO205 cells, we analysed another *BRAF*^*V600E*^ mutant colorectal cancer cell line, HT29, and observed a similar proportion of EdU positive cells during selumetinib treatment (Fig. S1B). Similarly, we observed replicating cells after treatment of COLO205 cells with the MEKi trametinib, showing that escape from G1 and entry to replication is not unique to selumetinib (Fig. S1C).

**Figure 2:**
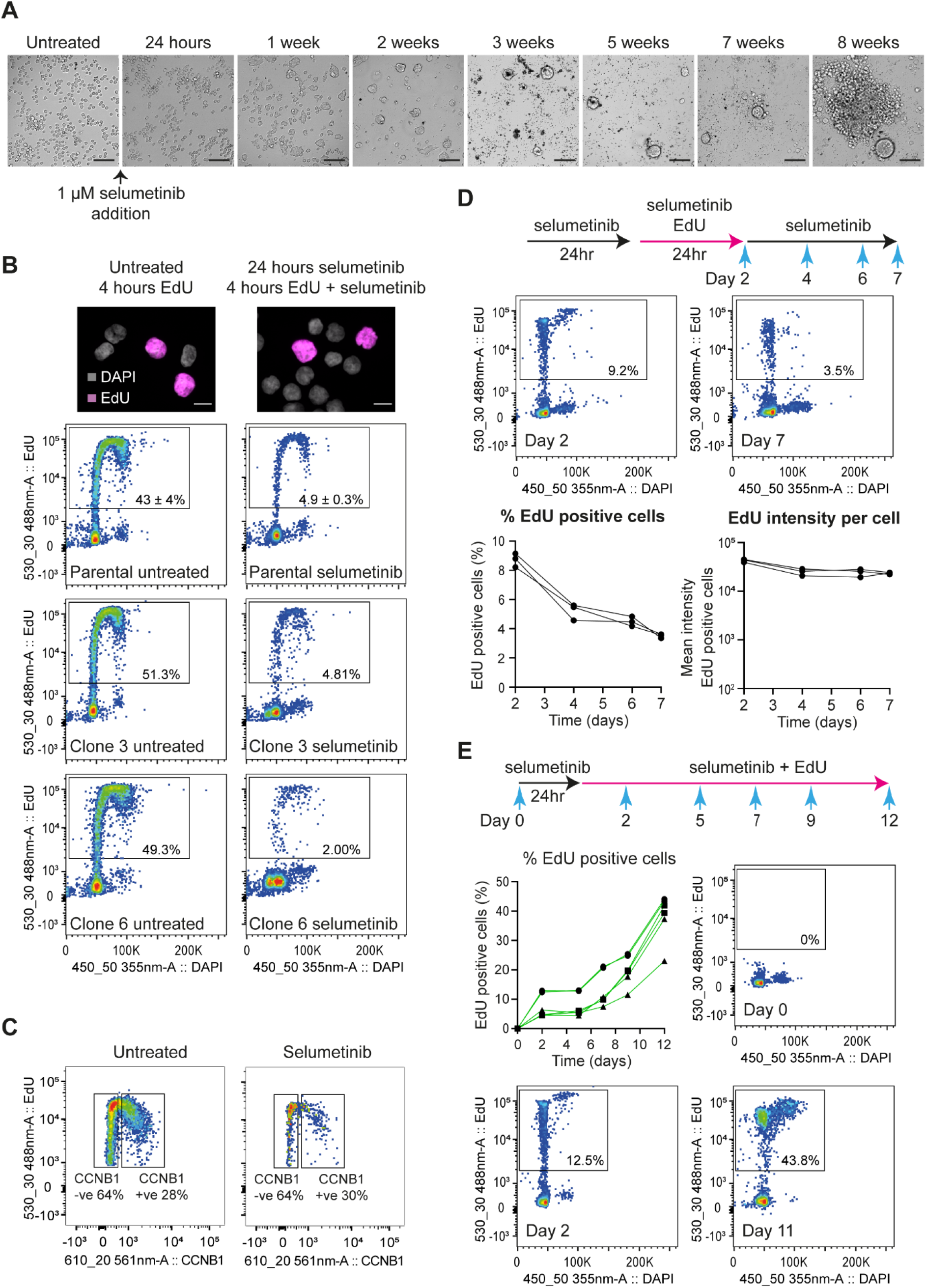
Replicating cells persist in long-term selumetinib-treated cell cultures. **A**: Representative brightfield images of COLO205 cells during extended treatment with 1 µM selumetinib (scale bars, 100 µm). **B**: EdU incorporation in COLO205 cells treated for 24 hours with 1 µM selumetinib before addition of 10 µM EdU for 4 hours. Representative images of EdU negative and positive cells (pink) co-stained with DAPI (grey) from selumetinib treated and control cells are shown at top (scale bars, 10 µm) and quantification of EdU positive cells in each population by flow cytometry shown below. Percent EdU positive cells are shown within the gates for each sample, figures for parental line are an average of 3 experiments with SD. **C**: Quantification by flow cytometry of CCNB1 negative and positive cells amongst the EdU positive cell population in untreated (left) and selumetinib treated (right) samples. COLO205 cells were treated with 1 µM selumetinib or DMSO only for 24 hours before addition of 10 µM EdU for 4 hours in the presence or absence of 1 µM selumetinib respectively. Cells were stained with EdU reaction cocktail and counter stained with CCNB1 primary antibody and Alexa Fluor 594 conjugated donkey anti-rabbit secondary antibody. Percent CCNB1 positive and negative cells in the EdU positive population are shown within the gates for each sample. **D:** COLO205 cells were treated with 1 µM selumetinib for 24 hours before addition of 2 µM EdU for 24 hours in the presence of 1 µM selumetinib, after which cells were rinsed in culture media and grown in the presence of 1 µM selumetinib only for up to 5 days. EdU incorporation was assayed at the indicated time points by flow cytometry. Results are mean of 2 independent replicates. Quantitation of EdU positive cells (left) and EdU intensity per cell (right), n=3 are shown in the bottom panel. **E:** Quantification of EdU positive cells by flow cytometry in COLO205 cells grown in the presence of selumetinib and EdU over the course of 11 days. COLO205 cells were treated for 24 hours with 1 µM selumetinib before addition of 2 µM EdU for 6 days, after which cells were rinsed with culture media then treated with 1 µM selumetinib and 2 µM EdU for a further 5 days. EdU incorporation was assayed at the indicated time points. Data for 6 independent replicates are shown.

After the 4-hour EdU pulse, ∼30% of EdU positive cells co-stained for the G2 marker Cyclin B1 (CCNB1) in both control and selumetinib treated populations (Fig. 2C), and DAPI incorporation of EdU/CCNB1 double-positive cells was consistent with 4n genome content (Fig. S1D), showing that cells progress through the cell cycle after escaping G1 arrest. The detection of cycling cells during extended selumetinib treatment could be explained by stochastic escape from G1 arrest or continued proliferation of a small subpopulation. To distinguish these, we performed an EdU pulse-chase experiment and observed that cells labelled during a 1-day EdU pulse did not increase in number or decrease in EdU intensity during the 5-day chase period, showing that cells labelled during the pulse did not re-enter the cell cycle (Fig. 2D). We then treated cells continuously for 11 days with EdU + selumetinib, during which time almost 50% of cells incorporated EdU (Fig. 2E). Together these experiments show that at least half the population can sporadically escape G1 arrest and undergo DNA replication during prolonged MEKi treatment.

### Gene expression during cell cycle progression in selumetinib

Suppression of ERK1/2 signalling is reported to downregulate DNA repair genes (30, 36-38), and indeed many DNA replication and repair genes were expressed at a significantly lower level in COLO205 cells after 24-48 hours of selumetinib treatment (Fig. S2A). DNA replication without normal expression of replication and repair genes is likely to be mutagenic, but it is unclear whether ERK remains inactive during sporadic re-entry to the cell cycle in drug, or whether these genes remain repressed since replicating cells in drug are too scarce to contribute to bulk mRNA-seq profiles.

High-content imaging for phosphorylated ERK1/2 (pERK1/2) showed that ERK is not reactivated in cells replicating in selumetinib, as pERK1/2 was equivalently reduced in EdU negative and positive populations under selumetinib treatment (Fig. 3A). To study gene expression in rare replicating cells, we developed a method for mRNA-seq after fixation, staining and sorting cells for intracellular markers (39), that we applied to CCNB1-positive G2 cells in selumetinib-treated and control populations (Fig. S2B). In accord with the pERK1/2 imaging data, CCNB1-positive cells in selumetinib did not re-express genes directly repressed by MEK inhibition (Fig. S2C) (40), nor display the known transcriptomic signature of MEK functional output (Fig. S2D) (41), so even if replication is initiated by transient ERK1/2 activation this has no lasting impact on the transcriptome. Comparing CCNB1 positive and negative cells in the presence and absence of selumetinib, 1681 genes were significantly and substantially (>4-fold) differentially expressed between conditions. These formed three hierarchical clusters: (i) genes expressed at a lower level under selumetinib treatment irrespective of cell cycle stage; (ii) genes expressed at a lower level in selumetinib-treated CCNB1 negative cells but expressed at normal levels in CCNB1 positive cells; (iii) genes expressed at a higher level under selumetinib treatment irrespective of cell cycle (Fig. 3B).

**Figure 3:**
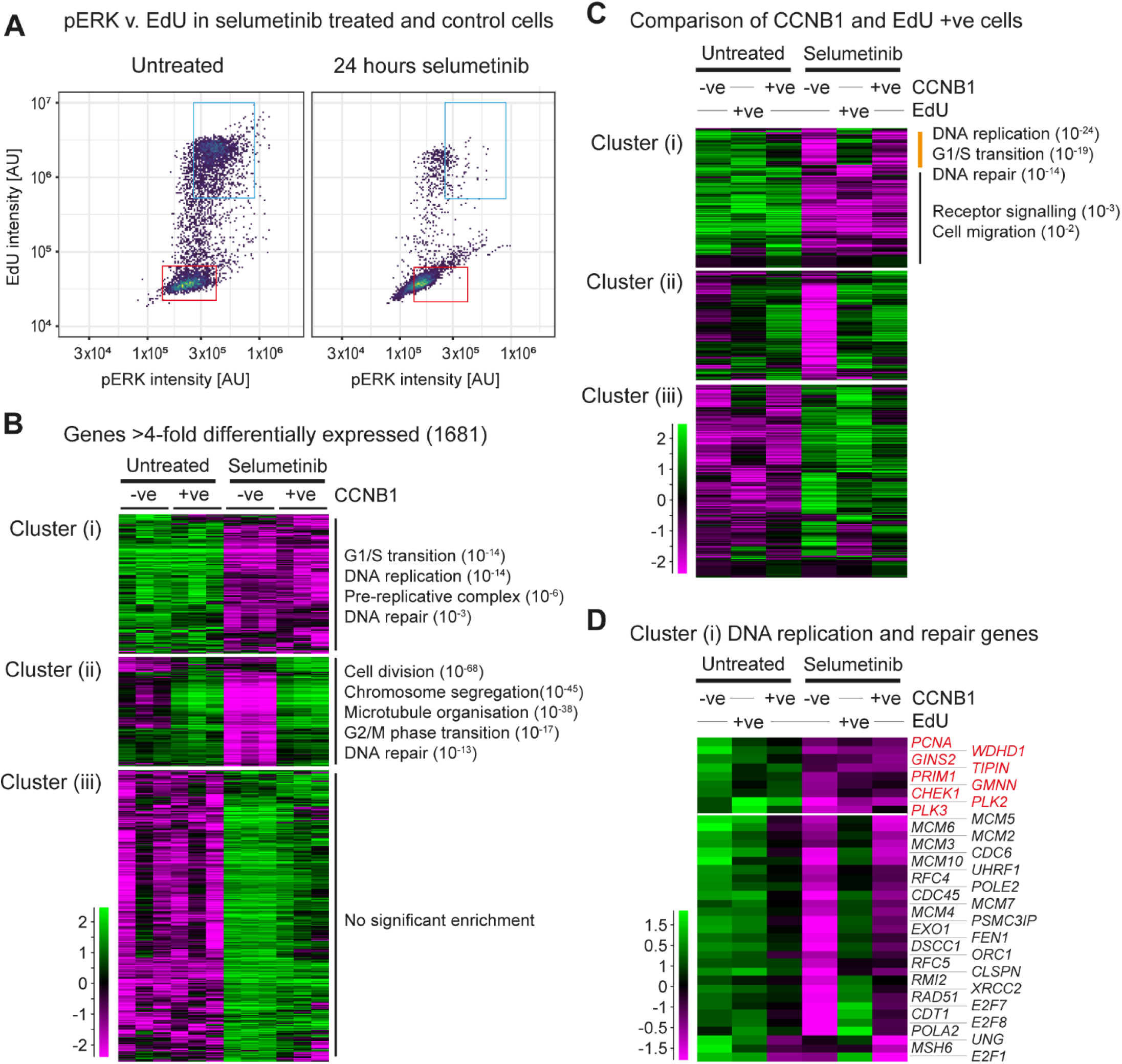
COLO205 cells replicating in selumetinib show defective gene expression. **A:** Quantification of pERK in EdU negative and positive cells. COLO205 cells were treated with 1 µM selumetinib or DMSO only for 24 hours before addition of 10 µM EdU for 4 hours in the presence of 1 µM selumetinib. Following EdU staining and immunofluorescence with pERK (T202/Y204) antibody, EdU incorporation and pERK levels in cells were determined by high content image analysis. EdU and pERK intensity in untreated (left) and selumetinib treated (right) cells normalised to control cells without addition of EdU or pERK primary antibody are shown. EdU negative and positive cells in individual plots are shown in red and blue rectangular gates respectively. **B:** Differential gene expression between CCNB1 negative and positive COLO205 cells, either untreated or after 24 hours selumetinib treatment. Cells from three biological replicates were fixed with glyoxal, then stained and sorted for CCNB1 followed by mRNA-seq library preparation. The 1681 genes shown are significantly (p<0.05 by DESeq2) and substantially (>4-fold) differentially expressed between at least one pair of the four categories shown. Genes were categorised into 3 primary behaviours by hierarchical clustering, and representative enriched GO categories (q<0.05) are shown (full GO analysis is presented in Table S1). **C:** Expression of gene clusters (i-iii) described in B in S/G2 cells. Cells were treated with 10 µM EdU for 4 hours prior to glyoxal fixation, followed by fluorophore conjugation, sorting and mRNA-seq. Cluster (i) splits into genes transiently upregulated in EdU +ve but not CCNB1 +ve cells (expressed in S but not G2), and genes equivalently expressed in EdU and CCNB1 +ve cells, separate GO analyses are shown for these sub-clusters, full GO analysis is presented in Table S2. **D**. Expression of cluster (i) genes associated with DNA replication in selumetinib treated and untreated populations sorted for CCNB1 or EdU, extracted from the dataset shown in C.

Most prominent in cluster (i) genes were GO terms relating to DNA replication, driven by transcripts encoding the entire MCM complex, replicative polymerase epsilon and alpha subunits, and other important replication proteins including PCNA, PRIM1, TIPIN and CLSPN, regulators such as PLK2, PLK3 and GMNN, and repair proteins RAD51 and EXO1. mRNA abundance for all these genes was low in both CCNB1 -ve and +ve populations under selumetinib treatment, whereas cluster (ii) transcripts are re-expressed to normal levels in CCNB1 +ve cells. GO analysis of this cluster reveals strong enrichments for chromosome segregation and also includes genes for DNA repair factors such as *BRCA2, BLM, GEN1* and *POLQ*, showing that chromosome segregation and DNA repair genes can be induced as required irrespective of ERK1/2 signalling. Cluster (iii) contained genes with a wide range of functions that were not significantly enriched for any GO category. To ensure that the behaviour of these gene sets is not unique to COLO205, we performed an equivalent experiment in HT29 cells, and observed that most genes in each of the three clusters were similarly affected by selumetinib, and that genes following the same expression patterns were enriched for similar GO categories (Fig. S2E).

These experiments show that key replication genes are mis-expressed in cells escaping selumetinib-induced G1 arrest, but profiling CCNB1-positive cells would miss a transient upregulation of transcripts during S phase. RNA recovered from EdU-treated cells is inevitably degraded during Click-labelling of EdU, but we were able to quantify transcript 3’ ends from EdU-positive cell samples (Fig. S2F). Reassuringly, cyclin mRNAs followed expected distributions - *CCNE1* and *CCNE2* were high in EdU-positive cells, *CCNB1* was high in CCNB1-positive cells, and *CCND1* was high in both (Fig. S2G). Across the three clusters defined above, the profiles of EdU-positive cells were similar to CCNB1-positive cells, but showed induction of some cluster (i) genes (Fig. 3C, orange bar). These genes were highly enriched for DNA replication and DNA repair categories and included the *MCM* genes, replicative polymerase subunits, *CLSPN, RAD51* and *EXO1*, showing that most key replication and repair genes are induced on sporadic entry to S phase during selumetinib treatment. However, the expression of *PCNA, WDHD1, PRIM1, TIPIN, PLK2, PLK3, CHEK1* and *GMNN* remained low across G1, S and G2 under MEKi treatment (Fig. 3D). Depletion of any of these genes decreases genome stability (42-47), providing support for the idea that DNA replication and chromosome segregation in MEKi-treated cells will be more error prone.

### Replication during MEK inhibition facilitates the emergence of drug resistance

Disrupted expression of *TIPIN, PLK2* or *PRIM1* increases replicative stress and reliance on ATR signalling (45, 48, 49) and indeed COLO205 cells show an increased sensitivity to ATRi in the presence of selumetinib (Fig. S3A). DNA replication during MEK inhibition should therefore carry an increased risk of *de novo* mutations such as the *BRAF* amplification. If *de novo* resistance mutations arise primarily because of errors during DNA replication or chromosome segregation in drug, then reducing the frequency of escape from G1 arrest should delay the emergence of drug resistance (Fig. 4A). The CCND1-CDK4/6 complex controls exit from G1, and MEKi and CDK4/6i combine to inhibit proliferation (50-52). This suggested that a low dose of CDK4/6i might enhance the selumetinib-mediated G1 arrest of selumetinib-sensitive parental cells, but not impair the proliferation of selumetinib-resistant cells in the presence of selumetinib (or of selumetinib-sensitive cells in the absence of selumetinib). By reducing the frequency of escape from the G1 arrest without impairing the outgrowth of any resistant cells that arise, we reasoned that we could quantify the effect of escape from G1 arrest in drug on formation of *de novo* resistance mutations.

**Figure 4:**
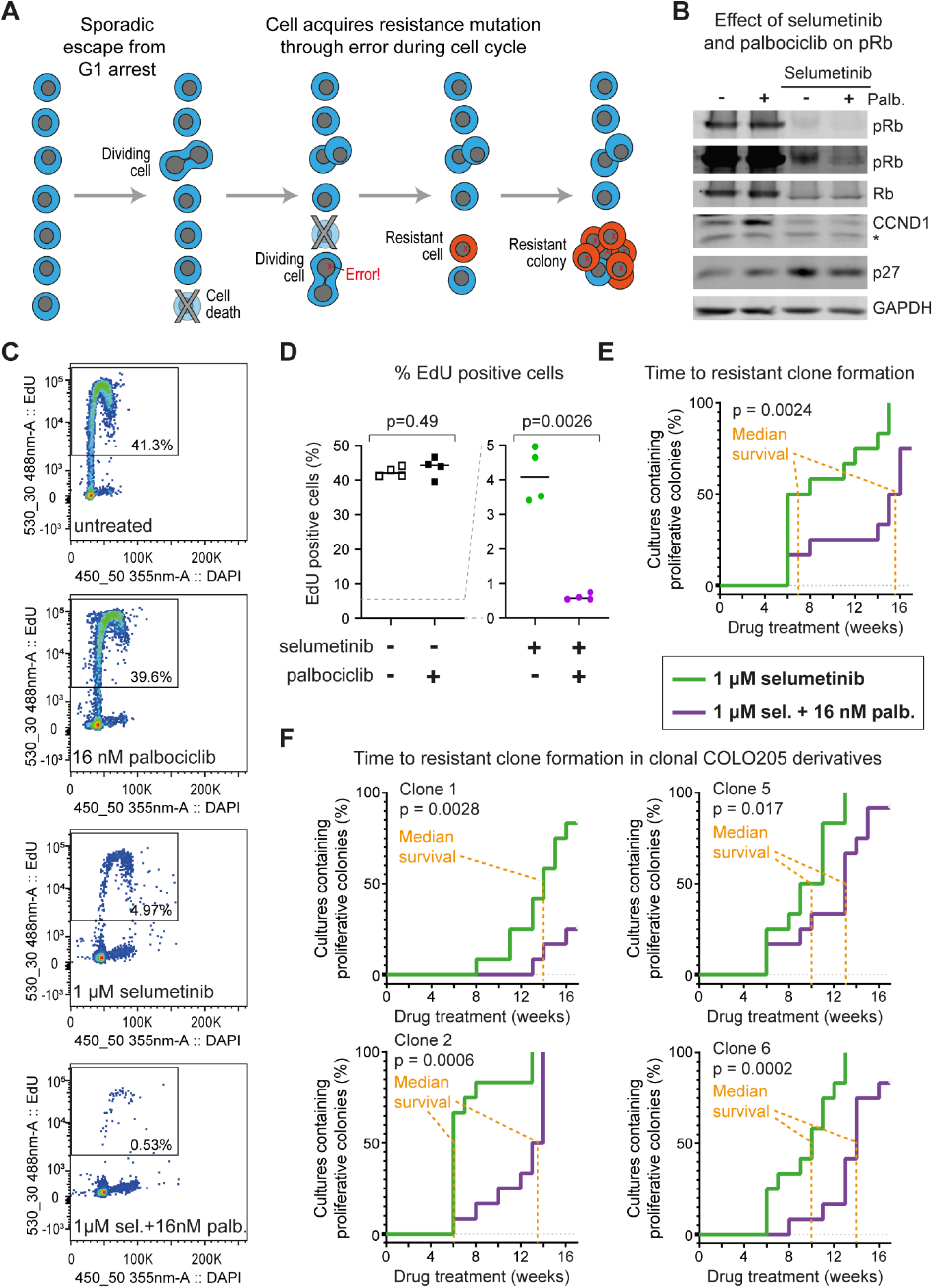
Suppressing DNA replication in selumetinib slows acquisition of resistance. **A**. Proposed mechanism for emergence of *de novo* resistance in drug. Selumetinib treated cells are arrested in G1 because of ERK1/2 pathway inhibition. Occasionally cells escape the G1 arrest and undergo a cell cycle, but this slow proliferation is offset by ongoing cell death and is not detectable in the bulk population. Occasional cell cycle events result in DNA replication or chromosome segregation errors that give rise to *de novo* mutations, some of which bestow drug resistance. This manifests as the sudden appearance of a rapidly proliferating colony after a long period of apparent stasis. **B**. Western blot analysis of COLO205 cells treated with 1 µM selumetinib in the presence (+) or absence (-) of 16 nM palbociclib for 24 hours and with the indicated antibodies. The pRB panels are shown at two different intensities to make the reduction of pRB levels in the combined selumetinib and palbociclib condition visible. Other panels show total Rb, CCND1, p27 (which is also part of the active CCND1-CDK4/6 complex), and GAPDH as a loading control. *indicates non-specific band **C**. Quantification of EdU positive cells by flow cytometry in COLO205 cells treated with palbociclib and/or selumetinib at the indicated concentrations for 24 hours before addition of 10 µM EdU for 4 hours. Percent EdU positive cells are shown within the gates for each sample. **D**. Quantification of EdU positive cells in **C**. n=4 (2 biological replicates each of parental COLO205 cells and single cell-derived clone 2), p values calculated by *t* test with Welch’s correction. **E**. Effect of combined treatment with selumetinib and palbociclib on time taken for the emergence of resistant clones in parental COLO205 cell line. Cells were cultured in media containing 1 µM selumetinib in the absence or presence of 16 nM palbociclib. Media and drug were replenished weekly, and the time taken for the appearance of first colony (≥50 cells) of proliferating cells in each well was recorded (in weeks). 12 independent replicates were performed under each condition, p values were calculated using the Mantel-Cox log-rank test on the principle that emergence of resistance can be represented as survival time of non-resistant cultures. **F**. Effect of combined treatment with 1 µM selumetinib and 16 nM palbociclib in single cell derivatives of COLO205 cells determined as in **E**. Data for 4 different single cell derivatives of COLO205 are shown.

We determined that treatment of COLO205 cells with the CDK4/6i palbociclib at 16 nM (∼ 10% IC_50_ (53)) did not reduce proliferation or colony formation in the absence of selumetinib (Fig. S3B, C), nor impair proliferation of selumetinib-resistant COLO205 cells in the presence of selumetinib (Fig. S3D). Rb phosphorylation was unaffected by 16 nM palbociclib alone, but residual Rb phosphorylation detected during selumetinib treatment was decreased by addition of 16 nM palbociclib (Fig. 4B). EdU incorporation assays confirmed that 16 nM palbociclib had no impact on entry to the cell cycle in the absence of selumetinib but reduced the fraction of EdU positive cells in the presence of selumetinib by 10-fold (Fig. 4C, 4D). Together these data show that 16 nM palbociclib alone does not cause G1 arrest, but substantially enhances the G1 arrest mediated by selumetinib.

We then compared the time taken for proliferating drug-resistant colonies to form in cultures treated with selumetinib alone or with a combination of selumetinib and 16 nM palbociclib in the parental COLO205 cell line and 4 single cell-derived clones. Resistant colonies formed in 86% of cultures, and average *BRAF* amplifications were equivalent for resistant cell lines derived under both conditions (Fig. S3E). However, addition of 16 nM palbociclib slowed the emergence of resistant clones substantially and significantly in all 5 cell lines, delaying the median time to resistance by three to eight weeks (Fig. 4E, F). We used a Cox Proportional-Hazards Model to quantify the overall effect of 16 nM palbociclib in combination with selumetinib compared to selumetinib alone, and found that 16 nM palbociclib reduced the risk of resistance by 78% with p=1.6×10^−11^. Differences between the COLO205 cell lines had no significant effect on palbociclib action (p=0.93), even though we observed a significant difference between the cell lines in acquisition of resistance in general (p=1.1×10^−10^). For example, clone 1 was slow to obtain resistance here as in Fig. 1D (p=5.5×10^6^) but took even longer to develop resistance under combined treatment with palbociclib.

This experiment shows that reducing the frequency of DNA replication during selumetinib treatment slows the formation resistant clones, and demonstrates that DNA replication in drug is the primary driver for the formation of *de novo BRAF* amplifications.

## Discussion

Here using BRAF^V600E^-mutant colorectal cancer cells we show that de novo *BRAF* amplifications arise in selumetinib-treated populations with remarkable efficiency. Cells under continuous selumetinib exposure stochastically escape G1 arrest and enter S phase but do so without inducing a subset of factors important for error-free DNA replication and chromosome segregation. Escape from G1 arrest is vulnerable to otherwise inert doses of CDK4/6i such that a MEKi+CDK4/6i combination suppresses DNA replication during selumetinib treatment, thereby retarding the formation of resistant clones.

Reduced expression of high-fidelity DNA replication and repair genes has been observed in drug-arrested cell populations (28, 30), which suggests that the sporadic entry of drug-treated cells into S phase may occur when replication factors are limited. However, our analysis of gene expression at specific cell cycle stages shows that DNA replication and repair genes are almost all induced as needed under MEKi treatment, so changes measured by bulk mRNA-seq largely reflect shifts in cell cycle distribution of the population. Nonetheless, a small number of genes important for accurate DNA replication and chromosome segregation are repressed across G1/S/G2 during MEKi treatment, which could underlie an increase in mutagenicity, and indeed our findings link the emergence of *de novo* resistance to cell cycle entry in drug.

One puzzling feature of drug tolerant persister (DTP) cells that survive for long periods in the presence of chemotherapeutics such as selumetinib is the sharp transition between drug tolerance and proliferation. The bulk population does not slowly re-acquire the ability to proliferate in drug; instead individual colonies of rapidly dividing cells suddenly appear after weeks or months of apparent stasis, requiring a marked return to proliferation in a very small number of cells (19-22). The mechanism we propose explains this property (Fig. 4A); occasional cell division events would not be noticeable in long-term drug treated cultures as these are offset by ongoing cell death (some of which may well arise through inappropriate entry to the cell cycle). However, if each replication event carries a risk of *de novo* gene amplification then each cell has a chance of acquiring the correct amplification to allow proliferation during a sporadic replication event. Gene amplifications arising in this manner would manifest as a sudden return of a single cell to proliferation, with an average time-to-resistance defined by the frequency of DNA replication events in drug and the extent to which drug treatment reduces the fidelity of replication.

Mutability under stress is well characterised in bacteria and has been repeatedly observed in yeast (54-57). However, it is hard to prove that such events result from defined programmes that have emerged through selective evolution, against the null hypothesis that mutagenesis is an emergent property of normal maintenance and proliferation systems becoming compromised under stress. We would therefore hesitate to label genome instability caused by under-expression of replication proteins as a mutagenic response, though our study provides strong support for the suggestion that non-genotoxic drug treatment can increase mutation rate and drive the emergence of resistance. Whether mutagenesis is intentional or not, our study and others addressing drug-induced mutation (28, 58) provide grounds for optimism that resistance to targeted chemotherapeutics is preventable, since mutational mechanisms that act during chemotherapy can be characterised and suppressed.

Overall, our study shows that pathways to *de novo* mutation can be mechanistically defined, and present vulnerabilities that can be specifically targeted to slow or stop the acquisition of drug resistance.

## Materials and Methods

Please see supplementary information for detailed methods.

### Cell culture and drug treatment

Cells were cultured in RPMI-1640 (COLO205) or McCoy’s 5A (HT29) media supplemented with 10% (v/v) foetal bovine serum, penicillin (100 U/mL), streptomycin (100 mg/mL) and 2 mM glutamine. Selumetinib and/or palbociclib (Selleck) resistant derivatives were generated by drug treatment with weekly replenishment of media and drug until proliferating colonies were observed. Cell line identity was validated using CellLineSleuth https://github.com/s-andrews/celllinesleuth.

### EdU staining and immunofluorescence for imaging

Cells were fixed in 4% formaldehyde and permeabilised in 0.5% triton X-100 before incubation in a reaction cocktail (43 µL Component D, 387 µL water, 20 µL CuSO_4_, 50 µL reaction buffer additive (43 µL 10x reaction buffer additive + 387 µL water) and 1.2 µL AlexaFluor 594 dye) (ThermoFisher Scientific) for 30 minutes at RT in dark and mounted in mounting medium with DAPI (Vector laboratories). For high-throughput imaging, cells cultured in 96-well plates (Perkin Elmer) were formaldehyde-fixed, permeabilised in ice-cold 100% methanol for 10 minutes at -20°C and labelled using an Alexa Fluor™ 647 HCS assay kit (ThermoFisher Scientific). Cells were blocked in 5% normal goat serum and 2% BSA for 1 hour, followed by incubation in primary antibody at 4°C overnight and secondary antibody for 1 hour at RT in dark. Cells were counterstained in DAPI and imaged using an INCell Analyser 6000 Microscope. Details of antibodies are provided in Table S5.

### EdU staining and immunolabelling for flow cytometry and sorting

EdU labelling was performed using a Click-iT™ EdU Alexa Fluor™ 488 Flow cytometry kit (ThermoFisher Scientific) following the manufacturer’s instructions, cells were counterstained in DAPI and analysed on a Fortessa (BD Biosciences) flow cytometer. Isolation of cells following CCNB1 staining was performed as described (39). For sorting by EdU, cells were incubated in a modified reaction cocktail (209 µL PBS, 5 µL CuSO_4_, 25 µL 1 M L-ascorbic acid (Sigma, A2174), 1.25 µL AlexaFluor 488 dye, 10 µL RNasin Plus) (ThermoFisher Scientific) and incubated on ice for 30 minutes in dark, prior to sorting.

### RNA extraction and mRNA-seq library preparation

RNA was extracted from cells using TRIreagent (Sigma) following manufacturer’s instructions and RNA integrity assessed using a Bioanalyzer 6000 pico chip (Agilent). mRNA-seq libraries were prepared using the NEBNext ultra (or ultra II) Directional RNA kit (NEB) with the NEBNext Poly(A) mRNA Magnetic Isolation Module.

### mRNA-seq data analysis

Reads were mapped to GRCh38 using HISAT2 v2.1.0 (59) by the Babraham Institute Bioinformatics Facility. Mapped data was imported into SeqMonk v1.47.0, DESeq2 analyses (60) performed within SeqMonk. Hierarchical clustering analysis was performed in SeqMonk, and GO analysis using GOrilla (61). Quoted p-values for GO analysis are FDR-corrected, for brevity only the order of magnitude is given. GEO accession: GSE168604.

### CNV microarray

DNA samples were processed by Cambridge Genomic Services (Cambridge University) for hybridisation onto cytoSNP 850K beadchips (Illumina) following the manufacturer’s instructions. Data were analysed with BlueFuse Multi software version 4.5 and the BlueFuse algorithm with default settings (10 contiguous markers for CNV and 500 contiguous markers for loss of heterozygosity (LOH)) and mapped to genome build 37. GEO accession: GSE168604.

### Protein extraction and Western blot

Preparation of cell lysates for SDS-PAGE and western blotting were performed as previously described (31). Total protein was separated in 10% SDS-PAGE gels for 4 hours at 75 V, transferred to methanol-activated immobilon-FL PVDF membranes (Merck Millipore) by wet transfer (0.2 M glycine, 25 mM Tris, 20% (v/v) methanol) at 20 V overnight. Membranes were blocked in a blocking buffer (5% milk in Tris buffered saline and Tween 20 (TBST) (5% (w/v) non-fat powdered milk, 10 mM Tris-HCl (pH 7.6), 150 mM NaCl, 0.1% (v/v) Tween-20) for 1 hour at RT followed by incubation with primary antibodies in 5% milk or 5% BSA in TBST overnight at 4°C and secondary antibodies (Table S5) for 1 hour at RT in dark. Bands were detected using a Li-Cor Odyssey Imaging System (LI-COR Biosciences).

### Colony formation assay

Cells (0.25×10^6^ cells/well) were seeded in 6-well plates and treated with 16 nM palbociclib or DMSO for 24 hours. 100 cells were then seeded per well in 6-well plates in the absence or presence of palbociclib and cultured with weekly media/drug replenishment for 21 days before crystal violet staining.

### Statistical analysis

Statistical tests were performed using GraphPad Prism v8.4.0, except the Cox Proportional-Hazards Model, which was implemented in RStudio v1.2.5033 https://github.com/segondsa/resistant-colonies.

## Acknowledgements

We greatly appreciate the assistance of Simon Andrews of the BI Bioinformatics facility for coding, Hanneke Okkenhaug of the BI Imaging facility for high-throughput imaging analysis, BI Flow, BI Next Generation Sequencing, BI Bioinformatics and BI Imaging facilities for service provision and experimental help, Fabiola Vacca for supporting experiments, Neesha Kara for figure preparation, Matthew Sale, Andrew Kidger, Kathy Balmanno and Fatima Santos for experimental assistance and useful discussions.

