## Supplementary figures for "DNA replication during acute MEK inhibition drives acquisition of resistance through amplification of the *BRAF* oncogene"

**Full Materials and Methods**

*Cell culture and drug treatment*

COLO205 and HT29 cell lines were provided by the laboratory of Dr. Simon J Cook (Babraham Institute). Cells were cultured in RPMI-1640 (COLO205) or McCoy's 5A (HT29) media supplemented with 10% (v/v) foetal bovine serum, penicillin (100 U/mL), streptomycin (100 mg/mL) and 2 mM glutamine at 37 °C in a humidified incubator with 5% (v/v) CO2. Selumetinib and/or palbociclib (Selleckchem) resistant derivatives were generated by culturing cells in indicated drug concentrations with media and drug replenished weekly until proliferating colonies formed in culture. To generate single-cell derivatives, cells ($5x10^6$ cells/mL) were incubated with 1 µg/mL DAPI (Sigma) and DAPI-negative cells sorted into 96-well plates containing media on a BD FACSAria III sorter (BD Biosciences).

Cell line identity was validated based on RNA-seq data generated in this work using Cell Line Sleuth, developed by Simon Andrews of the Babraham Institute Bioinformatics Facility (https://github.com/s-andrews/celllinesleuth).

*EdU staining and immunofluorescence for imaging*

Cells were fixed in 4% formaldehyde and permeabilised in 0.5% triton X-100 before incubation in a reaction cocktail (43 µL Component D, 387 µL water, 20 µL $CuSO_4$, 50 µL reaction buffer additive (43 µL 10x reaction buffer additive + 387 µL water) and 1.2 µL AlexaFluor 594 dye) (ThermoFisher Scientific) for 30 minutes at RT in dark and mounted in mounting medium with DAPI (Vector laboratories).

For high-throughput imaging, cells cultured in 96-well plates (Perkin Elmer) were formaldehyde-fixed, permeabilised in ice-cold 100% methanol for 10 minutes at -20°C and labelled using an Alexa Fluor™ 647 HCS assay kit (ThermoFisher Scientific). Cells were blocked in 5% normal goat serum and 2% BSA for 1 hour, followed by incubation in primary antibody at 4°C overnight and secondary antibody for 1 hour at RT in dark. Cells were counterstained in DAPI and imaged using an INCell Analyser 6000 Microscope. Details of antibodies are provided in Table S5.

*EdU staining and immunolabelling for flow cytometry*

EdU labelling was performed using a Click-iT™ EdU Alexa Fluor™ 488 Flow cytometry kit (ThermoFisher Scientific) following the manufacturer's instructions. EdU reaction mixture: 219 µL PBS, 5 µL $CuSO_4$, 25 µL 1x buffer additive and 1.25 µL Alexa Fluor dye. Cells were counterstained in DAPI and analysed on a Fortessa (BD Biosciences) flow cytometer.

To isolate cells by flow cytometry following CCNB1 staining, cells were processed as previously described (1). For sorting on EdU, cells were incubated in a modified reaction cocktail (209 µL PBS, 5 µL $CuSO_4$, 25 µL 1 M L-ascorbic acid (Sigma, A2174), 1.25 µL AlexaFluor 488 dye, 10 µL RNasin Plus) (ThermoFisher Scientific) and incubated on ice for 30 minutes in dark.

*RNA extraction and mRNA-seq library preparation*

RNA was extracted from cells using TRIreagent (Sigma) following manufacturer's instructions and RNA integrity assessed using a Bioanalyzer 6000 pico chip (Agilent). mRNA seq libraries were prepared using the NEBNext ultra (or ultra II) Directional RNA kit (NEB) with the NEBNext Poly(A) mRNA Magnetic Isolation Module and processed for sequencing as previously described in (1).

*mRNA-seq data analysis*

After adapter and quality trimming using Trim Galore (v0.5.0), RNAseq data was mapped to human genome GRCh38 using HISAT2 v2.1.0 (2) by the Babraham Institute Bioinformatics Facility. Mapped data was imported into SeqMonk v1.47.0 (https://www.bioinformatics.babraham.ac.uk/projects/seqmonk/) and normalised to total read count. DESeq2 analyses (3) was performed within SeqMonk using a p-value cut-off of 0.01, and significantly different genes were further filtered for genes with >4-fold difference in at least one comparison. RNA obtained from EdU-treated cells was of poor quality as the click reaction conditions cause some RNA degradation, so for comparisons involving these datasets (Figs. 3C, D) all datasets involved were treated as follows: 1) reads were filtered and all reads outside an annotated exon were discarded. 2) The GRCh38 annotation was parsed to yield annotations covering only the 3' 500 nucleotides only for high confidence transcripts using the code at https://github.com/s-andrews/three_prime_gtf, and opposite strand reads mapping to this annotation set were quantified. Restricting analysis to the 3' end reduces the impact of RNA degradation as the RNA-seq library preparation includes a poly(A) selection and therefore fragmented transcripts have a 3' end bias. 3) An enrichment normalisation to the 50$^{th}$ and 90$^{th}$ percentiles was performed in SeqMonk to match the distributions of the datasets, which minimises biases resulting from the differences in RNA quality. Hierarchical clustering analysis was performed using SeqMonk, and GO analysis of individual clusters performed using GOrilla (http://cbl-gorilla.cs.technion.ac.il/) (4, 5). Quoted p-values for GO analysis are FDR-corrected according to the Benjamini and Hochberg method (q-values from the GOrilla output), for brevity only the order of magnitude rather than the full q-value is given (6). GEO accession: GSE168604.

*DNA extraction and qPCR*

Genomic DNA from cells was extracted using a DNeasy Blood and Tissue kit (Qiagen), digested with *Eco*RI and purified using a QIAquick PCR purification kit (Qiagen). For each qPCR reaction, 4 µl DNA was mixed with 5 µL 2x Maxima SYBR mix, 0.2 µl each of forward and reverse primers (10 µM) (Table S6) and 0.6 µl water. Cycling condition: 95°C 10 minutes, then 40 cycles of (95°C 15 seconds, 60°C 1 min).

*CNV microarray*

DNA samples were processed by Cambridge Genomic Services (Cambridge University) for hybridisation onto cytoSNP 850K beadchips (Illumina) following the manufacturer's instructions. Data were analysed with BlueFuse Multi software version 4.5 and the BlueFuse algorithm with default settings (10 contiguous markers for CNV and 500 contiguous markers for loss of heterozygosity (LOH)) and mapped to genome build 37. The cluster and manifest files for processing CytoSNP-850K v1.1 were CytoSNP-850Kv1-1_iScan_C1_ClusterFile.egt and CytoSNP-850Kv1-1_iScan_C1.bpm respectively and CytoSNP-850Kv1-2_iScan_B1_ClusterFile.egt and CytoSNP-850Kv1-2_iScan_B3.bpm respectively for v1.2 beadchips. GEO accession: GSE168604.

*Protein extraction and Western blot*

Preparation of cell lysates for SDS-PAGE and western blotting were performed as previously described (7). Total protein was subjected to electrophoresis through a 10% SDS-PAGE gel for 4 hours at 75 V, transferred to methanol-activated immobilon-FL polyvinylidene difluoride membranes (Merck Millipore) by wet transfer (0.2 M glycine, 25 mM Tris, 20% (v/v) methanol) at 20 V overnight. Membranes were blocked in a blocking buffer (5% milk in Tris buffered saline and Tween 20 (TBST) (5% (w/v) non-fat powdered milk, 10 mM Tris-HCl (pH 7.6), 150 mM NaCl, 0.1% (v/v) Tween-20) for 1 hour at RT followed by incubation with primary antibodies in 5% milk or 5% BSA in TBST overnight at 4˚C and secondary antibodies (Table S5) for 1 hour at RT in dark. Bands were detected using a Li-Cor Odyssey Imaging System (LI-COR Biosciences).

*Colony formation assay*

Cells ($0.25x10^6$ cells/well) were seeded in 6-well plates and treated with 16 nM palbociclib or DMSO for 24 hours. Cells were harvested and 100 cells seeded per well in 6-well plates in culture media each in the absence or presence of palbociclib. Cells were incubated for 21 days, with media and drug replenished once a week. Colonies were stained with crystal violet (0.4% (w/v), Sigma) in 50% methanol.

*Statistical analysis*

Statistical tests were performed using GraphPad Prism v8.4.0, except the Cox Proportional-Hazards Model, which was implemented in RStudio v1.2.5033. Code is available on GitHub https://github.com/segondsa/resistant-colonies

### Supplementary Figures

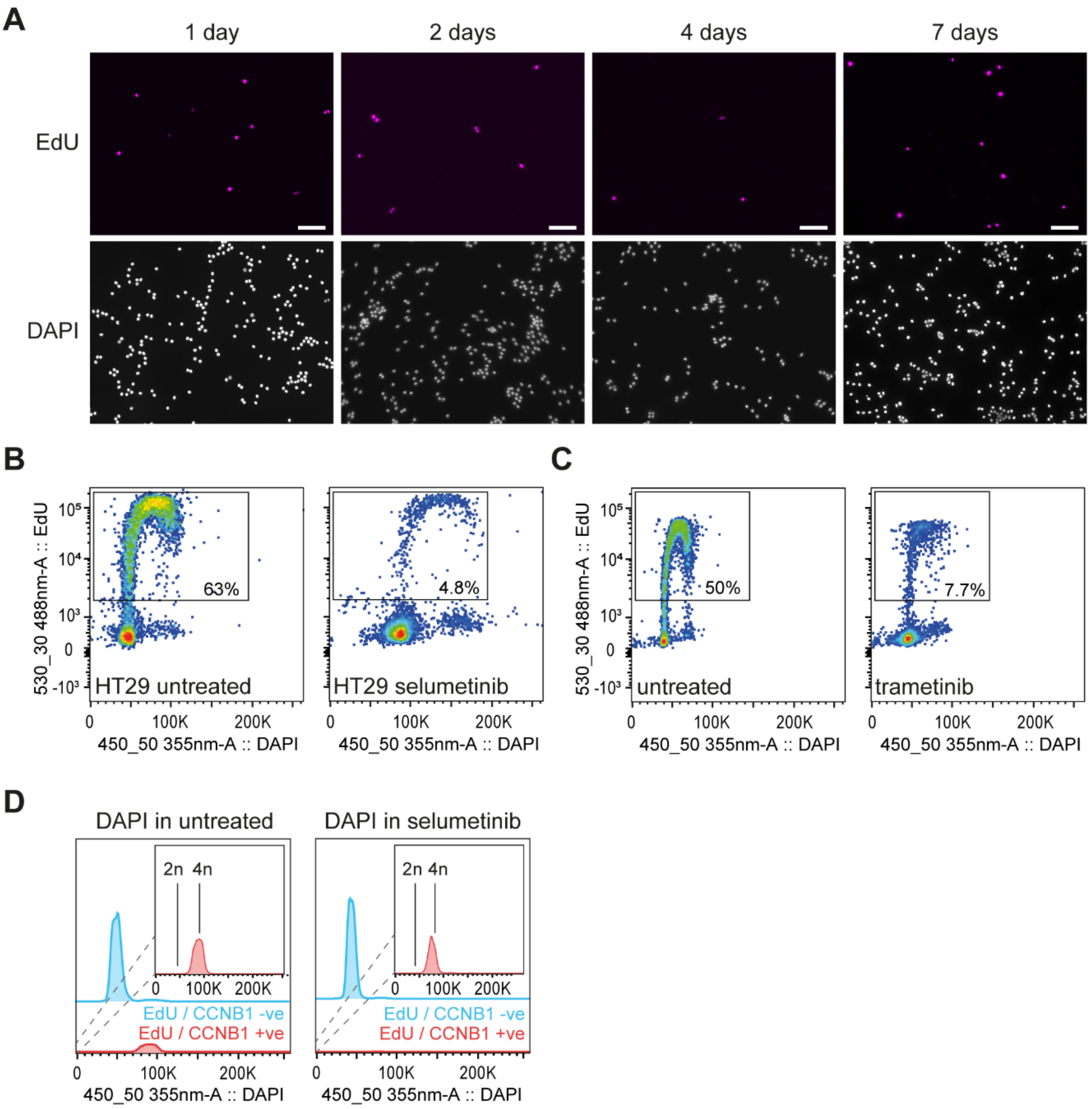

**Figure S1: Supplement to replicating cells persist in long-term Selumetinib-treated cell cultures**

**A.** EdU incorporation in COLO205 cells treated with 1 µM selumetinib for the indicated duration before addition of 10 µM EdU for 24 hours in the presence of selumetinib. EdU positive cells (pink) co-stained with DAPI (grey) from selumetinib treated cells are shown (scale bars, 100 µm).

**B.** Quantification of EdU positive cells by flow cytometry in HT29 cells treated with 1 µM selumetinib or DMSO only (untreated) for 24 hours before addition of 10 µM EdU for 4 hours. Individual plots in B and C show EdU incorporation and DAPI staining of DNA for untreated (left) and selumetinib treated cells (right), with rectangles to indicate gates used to quantify EdU positive and negative cells.

**C.** Quantification of EdU positive cells by flow cytometry in clonal single-cell derivative (clone 1) of COL0205 cells treated with 3 nM trametinib or DMSO only (untreated) for 24 hours before addition of 10 µM EdU for 24 hours in the presence of 3 nM trametinib.

**D.** Fluorescence histograms of DAPI intensities for EdU / CCNB1 double negatives and double positives in untreated (left) and 1 µM selumetinib treated (right) COLO205 cells. Cells were treated with 1 µM selumetinib or DMSO only for 24 hours before addition of 10 µM EdU for 4 hours. EdU incorporation, CCNB1 and DAPI incorporation were determined by flow cytometry. Inset plots show DAPI intensities of EdU-CCNB1 double positives re-scaled to make rare EdU / CCNB1 double positive signals visible.

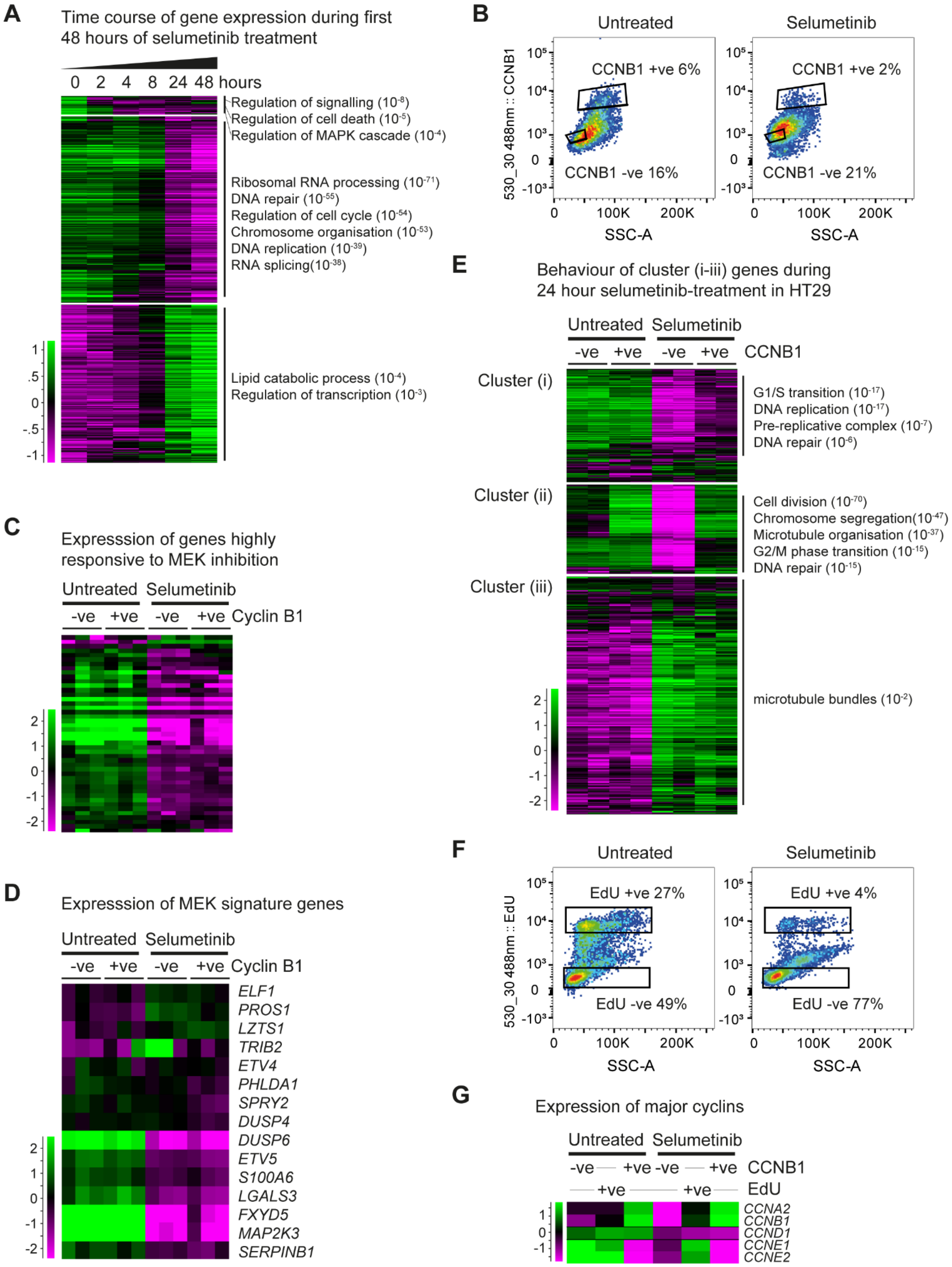
A
Time course of gene expression during first 48 hours of selumetinib treatment
0 2 4 8 24 48 hours
Regulation of signalling (10-8)
Regulation of cell death (10-5)
Regulation of MAPK cascade (10-4)
Ribosomal RNA processing (10-71)
DNA repair (10-55)
Regulation of cell cycle (10-54)
Chromosome organisation (10-53)
DNA replication (10-39)
RNA splicing(10-38)
Lipid catabolic process (10-4)
Regulation of transcription (10-3)
B
Untreated
Selumetinib
CCNB1 +ve 6%
CCNB1 -ve 16%
CCNB1 +ve 2%
CCNB1 -ve 21%
530_30 488nm :: CCNB1
SSC-A
C
Expresssion of genes highly responsive to MEK inhibition
Untreated Selumetinib
-ve +ve -ve +ve Cyclin B1
D
Expresssion of MEK signature genes
Untreated Selumetinib
-ve +ve -ve +ve Cyclin B1
ELF1
PROS1
LZTS1
TRIB2
ETV4
PHLDA1
SPRY2
DUSP4
DUSP6
ETV5
S100A6
LGALS3
FXYD5
MAP2K3
SERPINB1
E
Behaviour of cluster (i-iii) genes during 24 hour selumetinib-treatment in HT29
Untreated Selumetinib
-ve +ve -ve +ve CCNB1
Cluster (i)
G1/S transition (10-17)
DNA replication (10-17)
Pre-replicative complex (10-7)
DNA repair (10-6)
Cluster (ii)
Cell division (10-70)
Chromosome segregation(10-47)
Microtubule organisation (10-37)
G2/M phase transition (10-15)
DNA repair (10-15)
Cluster (iii)
microtubule bundles (10-2)
F
Untreated
Selumetinib
EdU +ve 27%
EdU -ve 49%
EdU +ve 4%
EdU -ve 77%
530_30 488nm :: EdU
SSC-A
G
Expression of major cyclins
Untreated Selumetinib
-ve +ve -ve +ve CCNB1
+ve +ve EdU
CCNA2
CCNB1
CCND1
CCNE1
CCNE2

**Figure S2: Supplement to gene expression analysis of replicating cells**

**A.** Gene expression across time during selumetinib treatment in COLO205 cells treated with 1 µM selumetinib for up to 48 hours, with cultures harvested at indicated times. The 5782 genes significantly differentially expressed ($p<0.05$ by DEseq2) between 2 replicates of 0 and 48 hour time points are shown. Genes were categorised into 3 primary behaviours by hierarchical clustering, and representative enriched GO categories ($q<0.05$) are shown (full GO analysis is presented in Table S3).

**B.** Flow cytometry density plots for COLO205 cells labelled with anti-CCNB1 primary antibody and donkey Alexa Fluor-488 conjugated secondary antibody and sorted using a BD FACSAria III sorter. Fluorescence thresholds for isolation of CCNB1 positive and negative cell fractions are shown. Gates were set based on a negative control staining without primary antibody, and the CCNB1 positive and negative sorting gates were set apart from each other to maximise sort purity.

**C:** Hierarchical clustering of gene set identified by Pratilas *et al*. (8) as significantly altered on MEK inhibition, showing relative expression in CCNB1 positive and negative fractions in the absence and presence of 1 µM selumetinib.

**D.** Hierarchical clustering of signature genes for MEK activity identified by Dry *et al.* (9) in CCNB1 positive and negative fractions in the absence and presence of 1 µM selumetinib.

**E.** Hierarchical clustering analysis of the three clusters of genes defined in Fig. 3B performed on HT29 cells either untreated or treated for 24 hours with 1 µM selumetinib and sorted for CCNB1, as in Fig. 3B. Full GO analysis is presented in Table S4.

**F.** Flow cytometry density plots for COLO205 cells stained for EdU and sorted using a BD FACSAria III sorter. Fluorescence thresholds for isolation of EdU positive and negative cell fractions are shown. Gates were set using the unstained negative control and the EdU positive and negative sorting gates were set apart from each other to maximise sort purity.

**G.** Expression of the mRNAs encoding major cyclins extracted from the data in Fig. 3C.

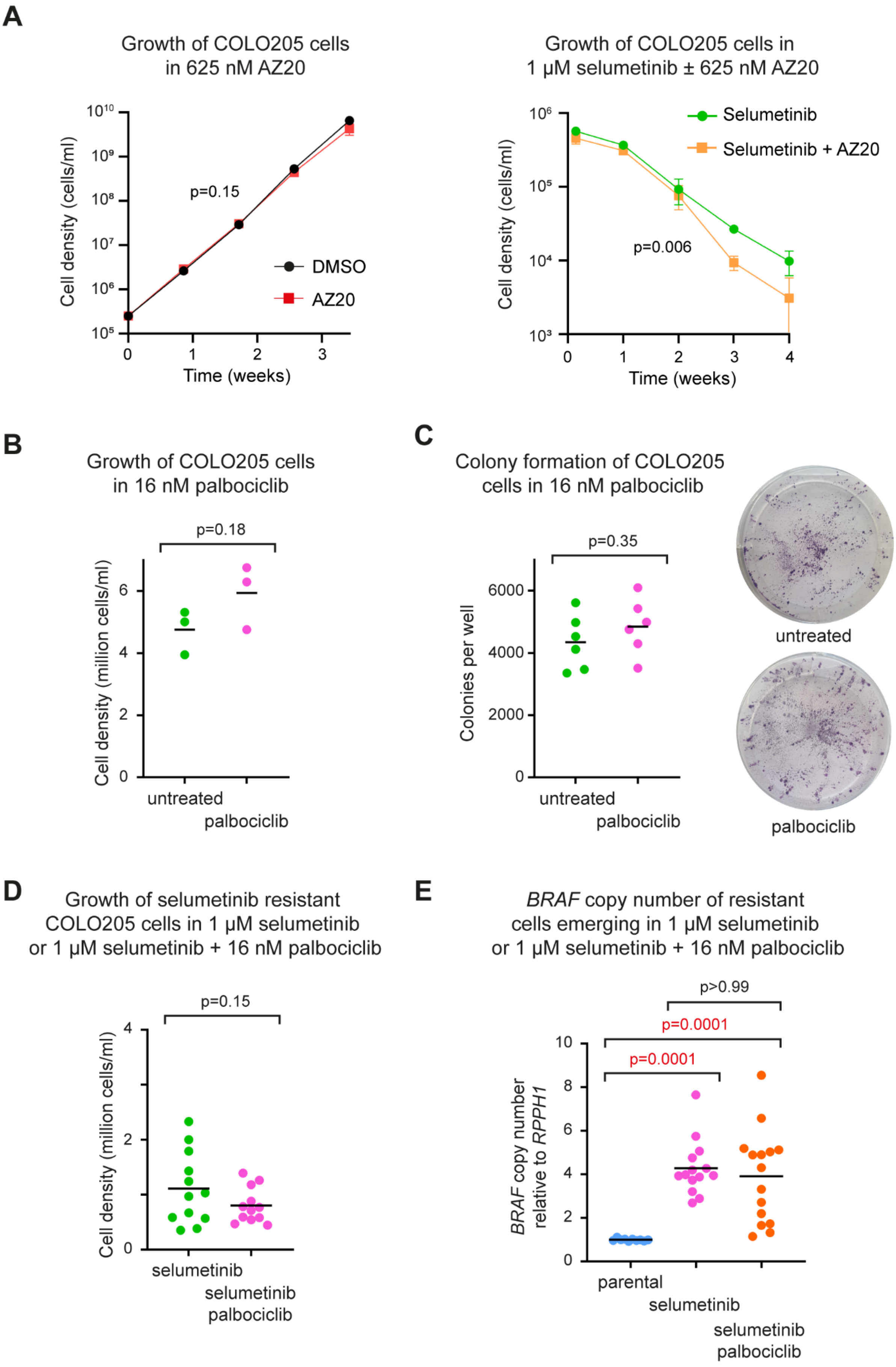
A
Growth of COLO205 cells in 625 nM AZ20
Cell density (cells/ml)
$10^{10}$
$10^9$
$10^8$
$10^7$
$10^6$
$10^5$
p=0.15
DMSO
AZ20
0
1
2
3
Time (weeks)
Growth of COLO205 cells in 1 µM selumetinib ± 625 nM AZ20
Selumetinib
Selumetinib + AZ20
Cell density (cells/ml)
$10^6$
$10^5$
$10^4$
$10^3$
p=0.006
0
1
2
3
4
Time (weeks)
B
Growth of COLO205 cells in 16 nM palbociclib
p=0.18
Cell density (million cells/ml)
0
2
4
6
untreated
palbociclib
C
Colony formation of COLO205 cells in 16 nM palbociclib
p=0.35
Colonies per well
0
2000
4000
6000
untreated
palbociclib
untreated
palbociclib
D
Growth of selumetinib resistant COLO205 cells in 1 µM selumetinib or 1 µM selumetinib + 16 nM palbociclib
p=0.15
Cell density (million cells/ml)
0
2
4
selumetinib
selumetinib palbociclib
E
BRAF copy number of resistant cells emerging in 1 µM selumetinib or 1 µM selumetinib + 16 nM palbociclib
p>0.99
p=0.0001
p=0.0001
BRAF copy number relative to RPPH1
0
2
4
6
8
10
parental
selumetinib
selumetinib palbociclib

**Figure S3: Supplement to suppressing DNA replication in selumetinib slows acquisition of resistance**

**A.** COL0205 cells were seeded at 0.25x10$^6$ cells/well in 6-well plates and treated with the indicated drugs (1 µM selumetinib and/or 625 nM AZ20) 24 hours later. Left: Every 6 days, cells were trypsinised, counted and 0.25x10$^6$ cells/well reseeded in media containing indicated drugs. Graph shows number of live cells at each time based on Trypan Blue staining, corrected for dilutions at each passage. Right: media and drug were replenished at weekly intervals without passaging, one well was harvested each week and live cells counted using Trypan Blue. All treatments were performed in parallel starting from the same cultures of cells, however we divided the data into two separate panels as the cells must be handled differently depending on whether they are proliferative (in the absence of selumetinib) or non-proliferative (in the presence of selumetinib). Error bars show SD, n=4 biological replicates for each condition, p values calculated by repeated measures two-way ANOVA.

**B.** COLO205 cells were seeded at 0.25x10$^6$ cells/well in 6-well plates and treated with 16 nM palbociclib or DMSO only for 24 hours, after which 100 cells from each condition were re-plated in media containing 16 nM palbociclib or DMSO only and cultured for 2 weeks with media and drug replenished weekly and counted using a Countess automated cell counter at the end of 2 weeks. p value calculated by t test, n=3 biological replicates.

**C.** COLO205 cells were treated and re-plated at 100 cells per well as in B to allow formation of colonies, then fixed and stained in 0.4% crystal violet in 50% methanol (representative images shown). Number of colonies was quantified, p value calculated by t test, n=6 biological replicates per condition.

**D**. Cell counts determined for selumetinib resistant cells derived from single-cell derivative of COLO205 cells (clone 2) as in B in the presence of 1 µM selumetinib. p value calculated by t test, n=12 biological replicates.

**E**: qPCR copy number analysis of *BRAF* relative to control gene *RPPH1* in parental and selumetinib-resistant cell lines derived in selumetinib alone or selumetinib + palbociclib. COLO205 cells were treated with 1 µM selumetinib in the presence and absence of 16 nM palbociclib, and media and drug changed weekly until colonies of proliferating cells were observed. Each sample was assayed in triplicate. p values were calculated by Kruskal-Wallis test (n=10 parental, 14 selumetinib, 15 selumetinib + palbociclib).
